# The Kamin Blocking Effect in Sign and Goal Trackers

**DOI:** 10.1101/2020.08.01.232553

**Authors:** Mayank Aggarwal, Jeffery R. Wickens

## Abstract

The discovery of the Kamin blocking effect suggested that surprise or prediction errors are necessary for associative learning. This suggestion led to the development of a new theoretical framework for associative learning relying on prediction error rather than just temporal contiguity between events. However, many recent studies have failed to replicate the blocking effect, questioning the central role of blocking in associative learning theory. Here, we test the expression of Kamin blocking in rats that either approach and interact with the conditioned cue (sign trackers) or approach and interact with the reward location (goal trackers) during appetitive classical conditioning. The behavioral task involved three phases: classical conditioning of a lever cue, conditioning of a compound of the lever cue plus an auditory cue, and testing response to presentation of the auditory cue in extinction. The results show that only sign trackers express the blocking effect. Thus, groups that include goal trackers are less likely to be able to replicate the blocking effect. Our findings support the idea that sign and goal tracking responses arise as a result of distinct parallel learning processes. Psychological theories of learning that incorporate these parallel learning processes and their interactions will provide a better framework for understanding the blocking effect and related associative learning phenomena.

## Introduction

Psychological principles underlying associative learning have undergone extensive experimental laboratory investigation using the classical conditioning paradigm introduced by Ivan Pavlov (Pavlov, 1927). In a typical classical conditioning experiment, the presentation of a cue is repeatedly followed the presentation of a reinforcer. After several pairings, the presentation of the cue comes to elicit a conditioned response that it did not elicit before. Early psychological theories of learning assumed that temporal contiguity between the cue and the reinforcer was sufficient for conditioning to occur. However, this assumption was called into question by the discovery of several behavioral phenomena in the latter half of the 20^th^ century, for example the Kamin blocking effect, latent inhibition and overshadowing.

Kamin blocking (Kamin, 1969a, 1969b) refers to the finding that conditioned responding to a cue is attenuated when it is paired with a reinforcer in the presence of another cue which has previously been conditioned using that reinforcer. The discovery of the blocking effect led to the suggestion that prediction error is necessary for learning to occur, stimulating the development of new associative learning theories that relied on prediction error rather than just temporal contiguity between events. Soon, the ability to explain the blocking effect became necessary for the validity of any theory of learning. However, many recent studies have failed to replicate the blocking effect, questioning the central status of blocking in associative learning theory (Maes et al., 2016; Maes et al., 2018; Soto, 2018). These failures also suggest a need to elucidate the precise experimental conditions necessary for the blocking effect.

The expression of blocking depends on what is learnt during appetitive classical conditioning. During appetitive classical conditioning two different types of conditioned responses can be acquired. Some subjects approach the conditioned cue and interact with it. This is called a sign tracking response (Boakes, 1977; Davey & Cleland, 1982; Hearst & Jenkins, 1974). Others approach and interact with the site of expected reward delivery when the conditioned cue is presented. This is called a goal tracking response (Boakes, 1977; Davey & Cleland, 1982; Farwell & Ayres, 1979; Holland, 1979).

Recent studies suggest that these differences in conditioned responding arise due to differences in the acquisition of incentive and predictive properties by the conditioned cue (Flagel et al., 2011; Robinson & Flagel, 2009). Incentive properties make the cue attractive, eliciting approach towards the cue, and are demonstrated by the general arousal elicited by the conditioned cue. Such arousal endows the conditioned cue with secondary reinforcing properties, which are only seen in those animals that develop a sign tracking conditioned response (Robinson & Flagel, 2009). This suggests that the conditioned cue develops incentive properties only in sign trackers. On the other hand, predictive properties acquired by the conditioned cue elicit reinforcer specific responses such as licking or gnawing, in both goal trackers and sign trackers (Davey & Cleland, 1982; Derman, Schneider, Juarez, & Delamater, 2018; Flagel et al., 2011). This suggests that conditioned cues develop predictive properties in both sign and goal trackers.

Given that a conditioned cue develops predictive properties in both sign and goal tracking animals, the prediction error explanation of the blocking effect suggests that both sign and goal tracking animals should express the blocking effect. However, the firing of midbrain dopamine neurons, which tracks the theoretical reward prediction error (RPE) signal (Hollerman & Schultz, 1998; Mirenowicz & Schultz, 1994; Schultz, Dayan, & Montague, 1997) suggests a different expression of the blocking effect in goal and sign trackers. Steinberg et al. (2013) suggested a causal role for the reduction in the dopamine response evoked by the expected reward in Kamin blocking. They found that optogenetic excitation of ventral tegmental area (VTA) dopamine neurons at the time of expected reward delivery attenuated blocking. Relating the dopamine RPE to sign and goal trackers, Flagel et al., (2011) found that the dopamine response evoked by the reward, when preceded by the cue being conditioned, declined only in sign trackers as classical conditioning progressed. In contrast, the reward evoked dopamine response did not decline in animals that developed a goal tracking conditioned response. Thus, the absence of the reduction in the dopamine response evoked by expected rewards in goal trackers leads to the hypothesis that goal trackers should not show the blocking effect.

Previous investigations into the expression of the Kamin blocking effect in animals that show a sign tracking or goal tracking conditioned response have produced mixed results. Derman et al., (2018) showed that both sign and goal tracking animals express the Kamin blocking effect. In contrast, Holland et al., (2014) found that goal trackers do not show the Kamin blocking effect. Other studies of Kamin blocking using pigeons showed that highly diffuse cues, which lead to the development of goal tracking conditioned responses, interfere with the development of sign tracking conditioned responses to localized cues (Blanchard & Honig, 1976; Khallad & Moore, 1996; Leyland & Mackintosh, 1978). However, diffuse cues do not support the sign tracking conditioned response because there is no discrete localized cue to direct responding towards, leaving no option but to goal track. Thus, goal tracking to a diffuse cue does not necessarily imply the acquisition of a goal tracking response in favor of sign tracking. When using diffuse cues, even though the conditioned response is goal tracking, the learning principles being followed may be the same as those underlying the development of a sign tracking conditioned response. This poses a problem in interpreting the results of experiments that use diffuse versus discrete cues to produce goal and sign tracking conditioned responding.

It is therefore important to test differences in learning between groups of animals that respond by either goal tracking or sign tracking to the same localized cue. That is, to use cues such that the cue identity is not variable across the goal and sign tracking groups. Both Derman et al., (2018) and Holland et al., (2014) used lever cues in phase 1 to establish sign tracking and, in a complimentary experiment, used auditory cues in phase 1 to establish goal tracking. In the experiment reported here, the expression of the Kamin blocking effect is compared in groups of animals that develop either a sign tracking or a goal tracking conditioned response to a lever cue paired with a food pellet reward.

## Material and Methods

The current paper presents a new analysis of data generated by previous experiments (Aggarwal, Akamine, Liu, & Wickens, 2020). Thus, the experimental procedures used here are the same as in Aggarwal et al. (2020), and are restated with minor modifications as follows.

## EXPERIMENTAL MODEL AND SUBJECT DETAILS

All animal experimental procedures were approved by the Committee for Care and Use of Animals at Okinawa Institute of Science and Technology. Male Long-Evans rats (Charles River Laboratories) aged 7-8 weeks on arrival were given food and water ad libitum and placed on a 12h light/dark schedule. Before starting habituation, they were food restricted to approximately 85% of their average body weight, and maintained at this weight throughout the behavioral experiments.

## METHOD DETAILS

### Behavioral Apparatus

Operant chambers were equipped with a food magazine, two levers, a house light, a white noise generator and a 4.5 kHz pure tone generator (Med Associates). The food magazine and the two levers were located on the same wall of the operant chamber, with one lever on either side of the food magazine. The two sound generators were placed on the wall opposite the food magazine.

### Habituation

Free exploration of the behavioral chamber was used to familiarize the rats to the experimental chamber. To allow free exploration, rats were placed in the chamber for 30 minutes on two consecutive days. No sound cues were presented, no food pellets were delivered, and no levers were present during free exploration. Prior reports suggest that such exposure is sufficient for familiarization to the experimental chamber (Bronstein, Neiman, Wolkoff, & Levine, 1974).

### Magazine training

After habituation, all rats underwent magazine training, which consisted of three sessions over three days. In each session there were 25 food reward deliveries (one sucrose pellet, 45mg) into the food magazine. The food deliveries were separated by a 60-80s variable interval (uniform random distribution).

### The Kamin blocking procedure

Only those rats that collected all the 25 food pellets on the last day of magazine training progressed to the Kamin blocking procedure. The blocking procedure consisted of three phases – a single cue conditioning phase, followed by a compound cue conditioning phase, and lastly an extinction test.

Phase 1 consisted of 12 single cue conditioning sessions over 12 days. Extension of the left and right levers for 5 seconds were used as the two cues. The retraction of one lever (L1, Fig. 1 was immediately followed by food delivery to the food tray. The other lever (L2, Fig. 1) was not followed by food reward. Each lever cue was presented 50 times in each session, with an inter-cue interval of 15-75s, and the cues were chosen randomly. Left and right levers were counterbalanced. The percentage of lever cue presentations that elicited a lever press response was used as the measure of conditioned responding (Davey, Oakley, & Cleland, 1981; Day, Roitman, Wightman, & Carelli, 2007).

**Figure 1:**
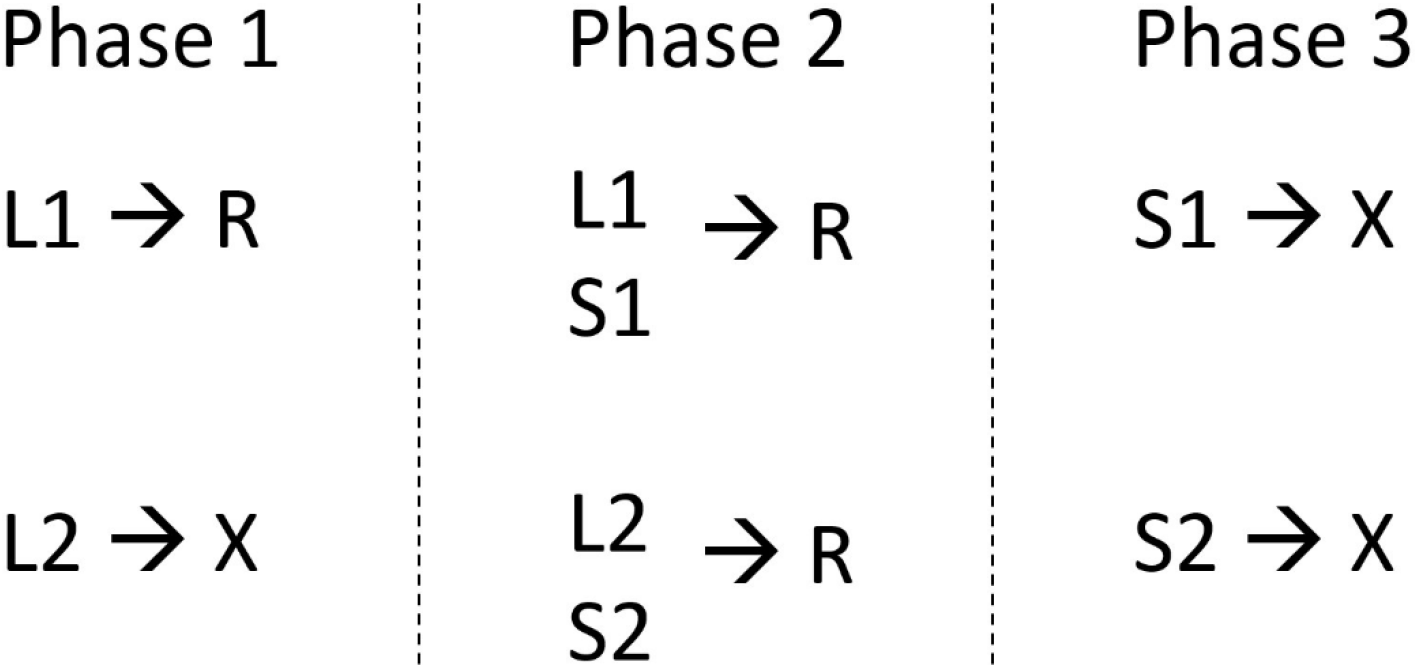
Schematic of the experimental design. During phase 1, the single cue conditioning phase, only one of the cues (L1 or L2) is followed by reward. During phase 2, the compound cue conditioning phase, compound cues (L1+S1 or L2+S2) are presented, and both compound cues are followed by reward. During phase 3, the extinction test, only the cues added in phase 2 (S1 and S2) are presented and no rewards are given.

Phase 2 of the Kamin blocking procedure consisted of 6 compound cue conditioning sessions over 6 days. Two compound cues were presented 25 times each in every session, with a variable inter-cue interval of 35-75s. One compound cue consisted of one of the lever cues used in phase 1 plus a simultaneous presentation of white noise (5s). The other compound cue consisted of the other lever cue plus simultaneous presentation of a pure tone (4.5 kHz, 5s). Cues were chosen randomly. The same food reward as used in phase 1 was delivered to the food tray immediately following the termination of each compound cue.

In phase 3, a single extinction session was used to assess the acquisition of conditioned responding (due to learning in phase 2) to separate presentations of the two sound cues. Each 5s sound cue was presented 25 times in this session, separated by a 35-75s variable interval. Sound cues were chosen randomly. Lever cues and rewards were not given during this session. The total duration of nose poking into the food magazine during the 5s presentation of the sound cues was used as the measure for conditioned responding. Nose poke duration during the 5s period immediately preceding each cue was used as the baseline response for that cue.

The inter-cue intervals were adjusted between phases to keep the session duration the same across phases, resulting in different inter-cue intervals in phase 1 than in phases 2 and 3.

## EXPERIMENTAL DESIGN AND STATISTICAL ANALYSIS

### Statistical Analysis

The development of a lever pressing conditioned response to the paired lever cue during phase 1 was assessed via a three way mixed ANOVA (Greenhouse-Geisser correction) on the lever press response measure, with session and cue as within subject factors, and group as the between subject factor. This was followed by an analysis of simple main effects to compare the difference in responding to the two lever cues within each group (L1 vs L2 in both the sign and goal tracking groups), and to compare the differences between the two groups in their response to each of the lever cues (sign vs goal tracking for both L1 and L2). Bonferroni adjustment was used to control for multiple comparisons.

To verify the development of a nose poking conditioned response to the lever cues during phase 1, a three way repeated measures ANOVA (Greenhouse-Geisser correction) was conducted on the relative response index, with session and cue as within subject factors, and group as the between subject factor. This was followed by an analysis of simple main effects to compare the difference in nose poke responding to the two lever cues within each group (L1 vs L2 in both the sign and goal tracking groups), and to compare the differences between the two groups in their response to each of the lever cues (sign vs goal tracking for both L1 and L2). Bonferroni adjustment was used to control for multiple comparisons.

Conditioned responding to the sound cues in phase 3 (extinction test) of the Kamin blocking procedure was assessed by comparing the average duration of nose poking during each of the sound cues (averaged over all 25 trials) with the average baseline nose poking preceding that cue. A two way mixed ANOVA (Greenhouse-Geisser correction) was conducted on the average nose poke duration measure, with cue as the within subject factor and group as the between subject factor. An analysis of simple main effects (Bonferroni adjustment) was done to compare the differences in responding elicited by the sound cues with their respective baselines within each group (S1 vs baseline_S1 and S2 vs baseline_S2 in both the sign and goal tracking groups).

We next wanted to compare inter-cue and inter-subject responding, because even if the subjects develop a conditioned response to both the cues, there could still be a difference in responding between the two cues. For this analysis, the response to each cue was normalized to each cue’s baseline response according to equation 1(Aggarwal et al., 2020).

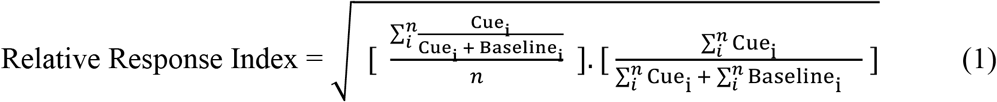

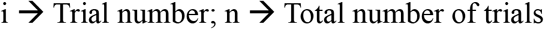

A two way mixed ANOVA was conducted on the relative response index with cue as the within subject factor and group as the between subject factor. This was followed by an analysis of simple main effects (Bonferroni adjustment) to compare the differences in conditioned responding to the two sound cues within each group (S1 vs S2 in both the sign and goal tracking groups) and the difference between the groups in their response to each cue (sign tracking vs goal tracking for both S1 and S2).

Responding to the compound cues during phase 2 was analyzed using two 3-way mixed ANOVAs, one on the lever press measure, and the other on the relative response index. Cue and session were the within subject factors with group being the between subject factor. An analysis of simple main effects (Bonferroni adjustment) was then used to compare differences in responding to the two compound cues within each group, and the differences between the groups in their response to each cue (sign tracking vs goal tracking for both L1 +S1 and L2+S2).

Statistical tests were considered significant if the probability of finding a false positive (type I error) was below 0.05 (alpha < 0.05). During the analysis of simple main effects, Bonferroni adjustment was used to control for multiple comparisons

### Inclusion Criteria

Animals had to satisfy either the sign tracking behavior criteria or the goal tracking behavior criteria to be included in the data analysis. Animals that did not pass either criteria, passed both the sign and goal tracking criteria, or developed a conditioned lever press response to the unpaired lever were excluded from data analysis.

Sign tracking behavior inclusion criteria: Aggarwal et al., (Aggarwal et al., 2020) used three quantitative criteria to assess the development of the lever pressing conditioned response to the levers, based on their preliminary studies. Here, we used the same criteria for inclusion on the basis of sign tracking behavior during phase 1 as follows. The rat had to pass any one of the three criteria for its response to the paired lever, and fail all three criteria for its response to the unpaired lever. The percentage of trials with a lever press response was used as the measure for responding to the levers. The following are the three quantitative criteria:

1.1) Above 40% response on the paired lever on one of the last three sessions.
1.2) Above 30% response on the paired lever on two of the last six sessions.
1.3) Above 20% response on the paired lever on three of the last six sessions.

Goal tracking behavior inclusion criteria: Criteria were developed to assess the development of a nose poke conditioned response to the paired lever, and the absence of such a response to the unpaired lever. The rats had to satisfy the following two goal tracking behavioral criteria to be included in the data analysis as goal trackers.

1. The rats had to satisfy one of the following two conditions assessing conditioned responding to the paired lever.

a. Relative response index for the paired lever above 0.65 on the last two sessions and above 0.7 on at least one of the last two sessions.
b. Relative response index for the paired lever above 0.7 on the last session and on at least three of the last four sessions.
2. The rats had to satisfy one of the following two conditions. This requirement eliminated those who had developed a conditioned response to the unpaired lever.

a. Relative response index for the unpaired lever below 0.7 on the last four sessions, below 0.65 on at least two of the last four sessions, and below 0.6 on at least one of the last four sessions.
b. Relative response index for the unpaired lever below 0.7 on the last session and below 0.6 on three of the last four sessions.

Out of 59 rats, 26 rats met the sign tracking criteria and 18 rats met the goal tracking criteria. Sixteen rats did not meet either criterion and three rats developed a lever press conditioned response to the unpaired lever. These 19 rats, and four rats that met both criteria were excluded from data analysis. Thus, the sign tracking group consisted of 22 rats and the goal tracking group consisted of 14 rats.

As the classification criteria of goal tracking is somewhat arbitrary, we also examined the effect of using two alternative classifications of goal tracking, in which, for inclusion as goal trackers, the average of the relative response index of the last three (first alternative classification) or last four (second alternative classification) sessions of phase 1 had to be above 0.65 for the paired lever and below 0.65 for the unpaired lever, with a minimum difference of 0.1 between the two levers. Both these alternative classifications also lead to a similar classification of 22 sign trackers and 14 goal trackers, and 4 rats common to both groups.

## Results

Classical conditioning using lever cues paired with a food pellet reward resulted in the development of sign tracking in some animals, and goal tracking in others. Figure 2 shows the gradual development of a conditioned response to the lever cue paired with the food reward in both the sign trackers and goal trackers. The sign trackers lever pressed when the lever paired with the food reward was presented and did not lever press when the unpaired lever was presented during phase 1. The goal trackers did not develop a lever pressing conditioned response (Fig 2A) but developed a nose poking conditioned response into the food magazine (Fig 2B).

**Figure 2:**
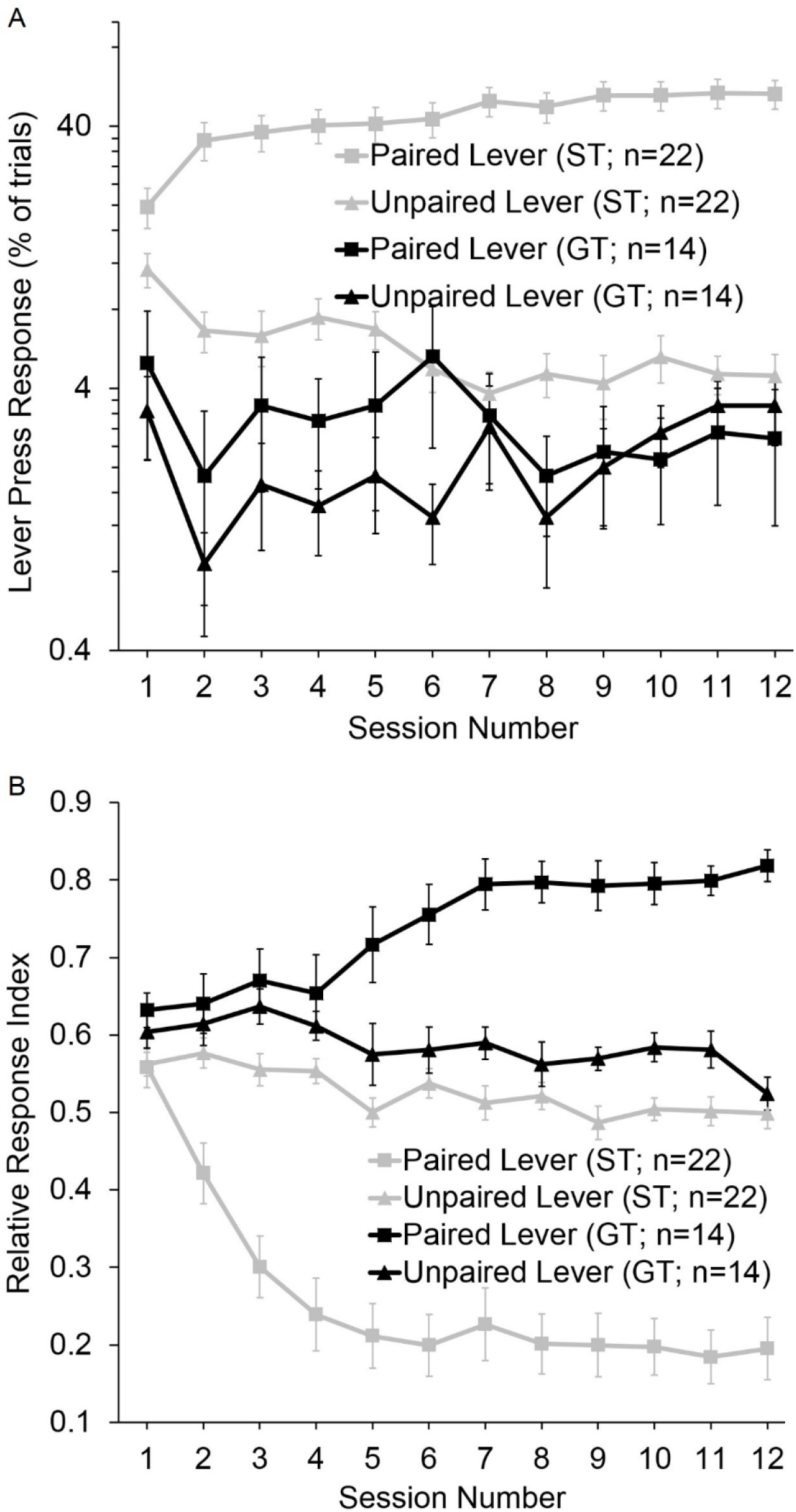
Conditioned responses of the sign tracking and goal tracking groups during phase 1. (A) Lever press responses of sign trackers and goal trackers during phase 1 of the Kamin Blocking Paradigm. (B) Nose poke responses of sign trackers and goal trackers to lever cues during phase 1. Nose poke responses are normalized to their baselines (response during the 5s immediately preceding the respective lever cue) according to equation 1. Data are presented as mean ± SEM.

Statistical analysis showed that the differences in behavioral patterns apparent in figure 2 were statistically significant. A three way mixed ANOVA on the lever press response measure, using the Greenhouse-Geisser adjustment, showed a significant effect of lever cue (F=51.526; p<0.001), a significant effect of group (F=110.147; p<0.001), and a significant cue x session (F=3.449; p=0.003), cue x group (F=57.825; p<0.001) and cue x session x group (F=5.516; p<0.001) interaction. A simple main effects comparison using the Bonferroni adjustment showed a significant effect of the lever cue in only the sign trackers (F=140.477; p<0.001) but not the goal trackers (F=0.074; p=0.787), showing the development of a lever pressing conditioned response only in the sign trackers. This means that only the sign trackers developed a lever pressing conditioned response in response to the lever cue paired with the reward, while the goal trackers did not develop a lever press response to either of the lever cues.

Further statistical analysis was necessary to verify the development of a nose poking conditioned response in the goal tracking group. A three way repeated measures ANOVA (Greenhouse-Geissier correction) on the relative response index (normalized nose poke response measure; Equation 1) for the goal trackers and sign trackers showed a significant main effect of cue (F=6.608; p=0.015), session (F=5.317; p<0.001) and group (F=82.323; p<0.001). In addition, all interactions were also significant – cue x group (F=92.064; p<0.001), cue x session (F=3.728; p=0.001), group x session (F=17.598; p<0.001), and cue x session x group (F=16.062; p<0.001). A simple main effects comparison (Bonferroni adjustment) showed a significant effect of the lever cue in the goal trackers (F=20.186; p<0.001). This means that the goal trackers developed a nose poking conditioned response only to the paired lever. There was a significant effect of the lever cue in sign trackers as well (F=95.144; p<0.001). This occurred because the nose poking response during L1 was suppressed below baseline in the sign trackers, as they were interacting with the lever during L1. Thus, the goal trackers developed a nose poking conditioned response in response to the lever cue paired with the reward, while the sign trackers suppressed their nose poke responses to the presentation of L1. There was also a significant effect of session on the nose poke response to L1 in both the goal tracking (F=3.265; p=0.007) and sign tracking groups (F=18.911; p<0.001). There was no significant effect of session on nose poke responses to L2 in the sign (F=1.713; p=0.131) or goal (F=1.194; p=0.342) tracking groups. This means that, over the course of multiple conditioning sessions, goal trackers developed a nose poke conditioned response and sign trackers developed a suppression in their nose poke response only to that lever which was paired with the food reward.

Both the groups then underwent compound cue conditioning (phase 2). When their response to the presentation of the sound cues alone was tested in extinction (phase 3), the sign trackers increased their nose poke duration from baseline nose poke duration only during the sound that was given as a compound along with the unpaired lever in phase 2 (S2) (t=5.463; p<0.001; two way mixed ANOVA – simple main effects – Bonferroni). The sign trackers did not increase their nose poke duration from baseline during the sound cue that was given as a compound along with the paired lever in phase 2 (S1) (t=1.596; p=0.728) (Fig. 3A). These findings show that the sign trackers responded to S2 but not to S1, producing the Kamin blocking effect. The goal trackers responded to both S1 (t=5.095; p<0.001) and S2 (t=5.277; p<0.001) (Fig. 3A), suggesting that the Kamin blocking effect was attenuated in this group. In addition, there was a significant difference in the average nose poke duration during S1 between the two groups (F=5.015; p=0.032) but not in the average nose poke duration during S2 (F=0.402; p=0.530) nor during the baseline period of S1 (F=0.484; p=0.491) and S2 (F=0.014; p=0.908). This suggests that the two groups differed significantly only in their nose poke response to S1, with the goal tracking group showing significantly longer nose poking than the sign tracking group during S1.

**Figure 3:**
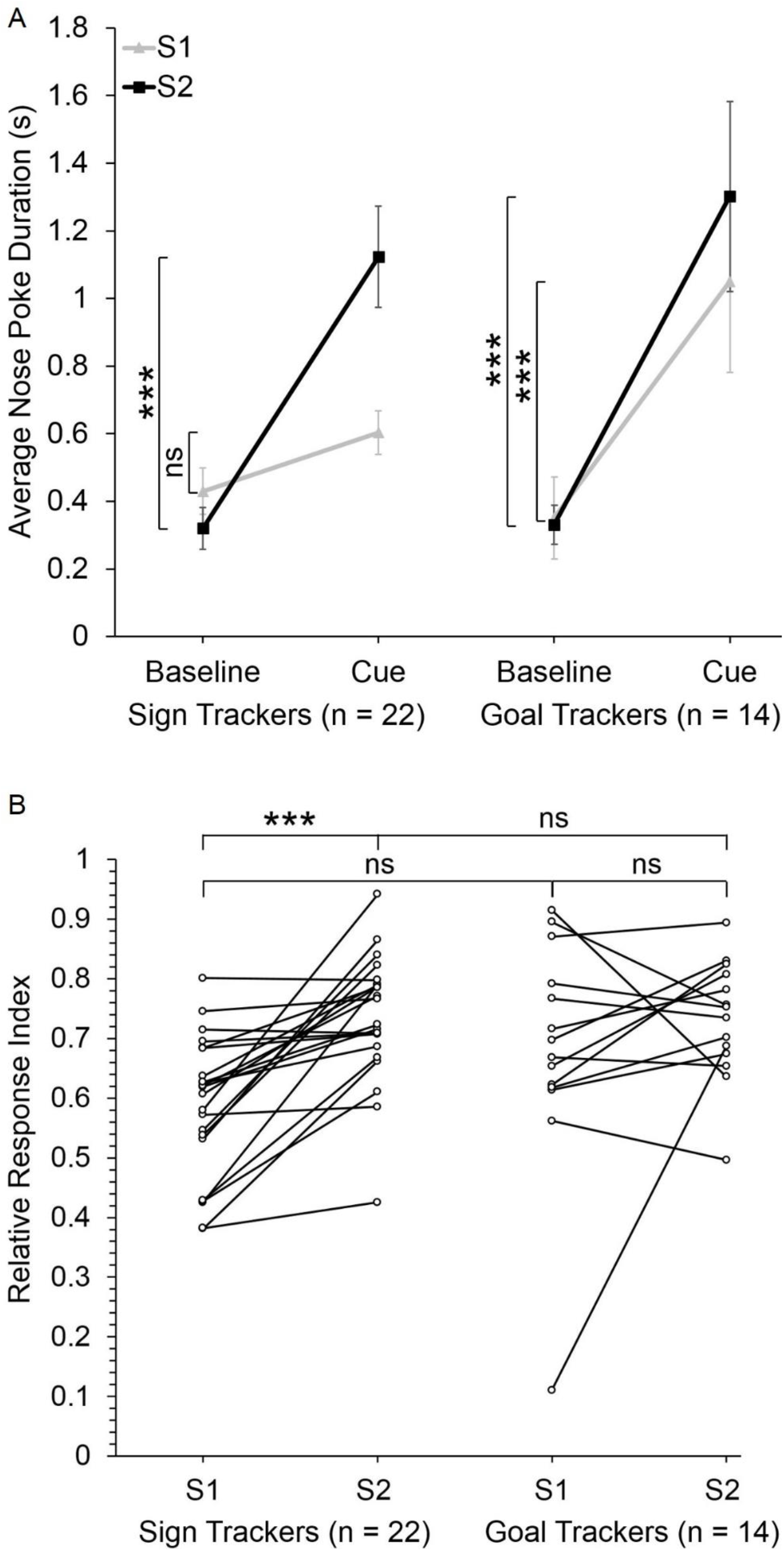
Kamin blocking effect is robust in sign trackers but attenuated in goal trackers. (A) Nose poke responses for sign and goal trackers when the sound cues are presented in phase 3. Baseline response is measured during the 5s immediately preceding the respective sound cue. Cue response is measured during the cue, which lasts for 5s. (B) Responses to the sound cues normalized (equation 1) to their respective baselines. Data are presented as mean ± SEM.

In agreement with the foregoing comparison of the raw nose poke scores, when the responses to the sound cues were normalized to their respective baselines (Equation 1), statistical tests showed an attenuation of the Kamin blocking effect in the goal tracking group. A two way mixed ANOVA showed a significant main effect of cue (F=14.164; p=0.001). There was no significant effect of group (F=1.534; p=0.224) or interaction between cue and group (F=3.306; p=0.078). A simple main effects comparison using the Bonferroni adjustment showed that the sign trackers had a significantly higher relative response index for S2 than for S1 (F=20.029; p<0.001). The goal trackers did not show a difference in their relative response index for S1 and S2 (F=1.548; p=0.222) during phase 3 (Fig. 3B). This means that the sign tracking group showed a robust Kamin blocking effect. The measure of the Kamin blocking effect did not reach significance in the goal tracking group, implying that the effect was attenuated in this group of animals. There was no significant difference in the relative response index of the two groups for S1 (F=3.16; p=0.084) and S2 (F=0.011; p=0.917) (Fig. 3B).

These results show that animals that develop a sign tracking conditioned response to the lever cue paired with the food pellet reward during phase 1 express the Kamin blocking effect. In contrast, the blocking effect is attenuated in animals that develop a goal tracking conditioned response to the paired lever cue.

It is important to note here that the pattern of responding to S1 and S2 in the sign and goal tracking groups observed in phase 3 could occur either due to differences in nose poking during S1 and S2 or due to differences in nose poking during the baseline period of S1 and S2. The baseline period is the 5s immediately preceding the presentation of a sound cue. Nose poking during the 5s immediately preceding S1 was not significantly different from nose poking before S2 in the sign tracking (t=2.037; p=0.297) and goal tracking groups (t=0.294; p=1.0). There was also no significant difference between the two groups in their baseline before S1 (t=0.693; p=0.491) or S2 (t=0.12; p=0.908). These results show that the pattern of conditioned responding to S1 and S2 observed in phase 3 in the two groups is not due to differences in baseline nose poking.

To check whether the pattern of responding observed in phase 3 was already present in phase 2, a 3-way mixed ANOVA was conducted on the relative response index for the two compound cues during phase 2 (Fig. 4B). A simple main effects comparison using Bonferroni adjustment showed that there was a significant difference in the relative response index for the two compound cues in the sign trackers (F=57.898; p<0.001) but not in the goal trackers (F=2.706; p=0.109). There was also a significant difference between the two groups in their relative response index for L1+S1 (F=143.624; p<0.001), but not for L2+S2 (F=0.07; p=0.793). These differences occurred because nose poke responding during L1+S1 in the sign trackers was suppressed below baseline, just as it was suppressed on L1 presentation during phase 1 in this group. This suppression in nose poking was not observed in phase 3 in the sign tracking group. Thus, the significant differences observed in phase 2 do not necessarily foreshadow the significant differences observed in phase 3.

**Figure 4:**
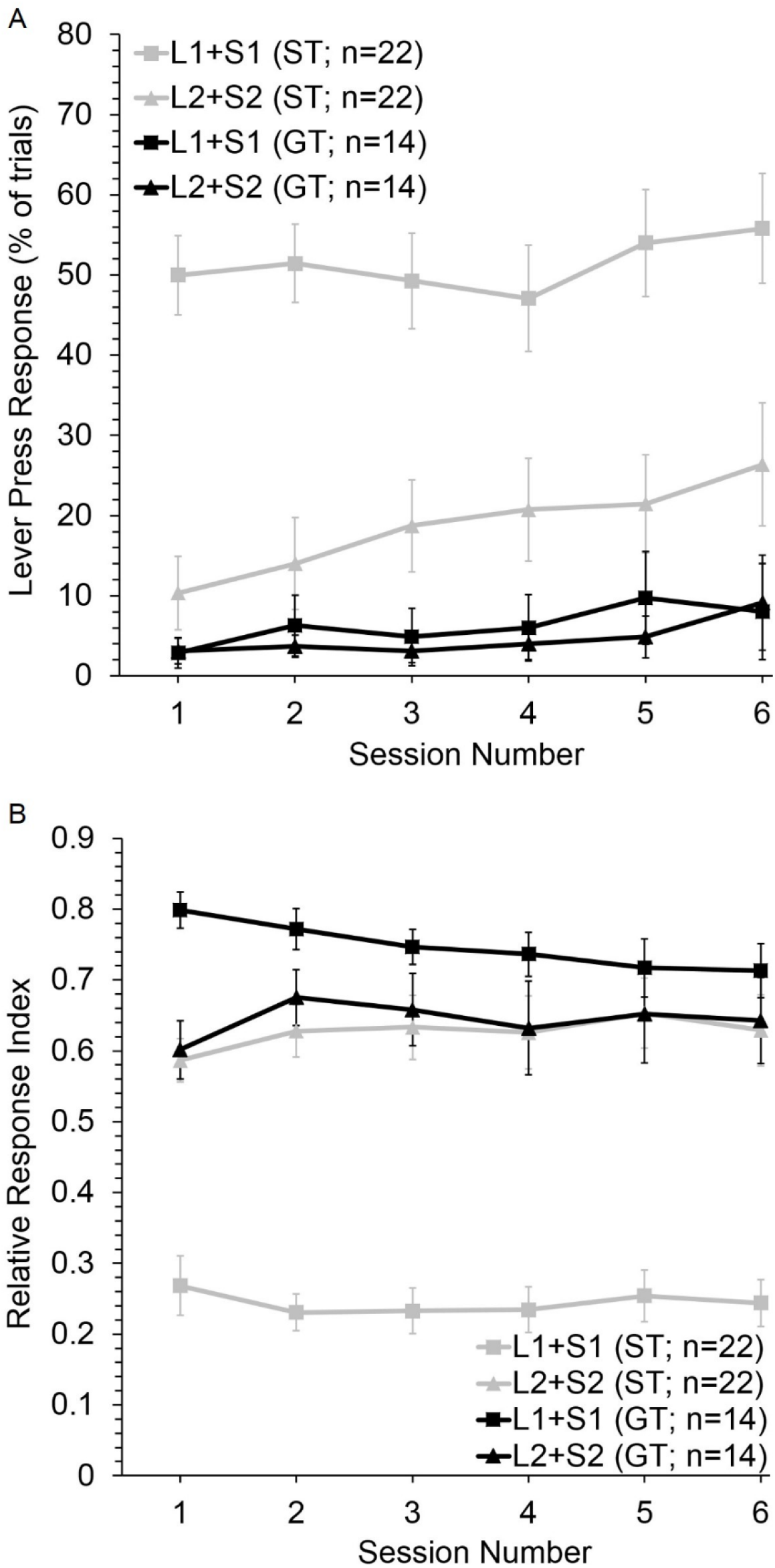
Lever and nose poke responses of the sign tracking and goal tracking groups during phase 2. (A) Lever press responses of sign trackers and goal trackers to the two compound cues, L1+S1 and L2+S2, during phase 2 of the Kamin Blocking Paradigm. (B) Nose poke responses of sign trackers and goal trackers to the two compound cues lever cues during phase 2. Nose poke responses are normalized to their baselines (response during the 5s immediately preceding the respective lever cue) according to equation 1. Data are presented as mean ± SEM.

The observed suppression in nose poking to L1+S1 in the sign tracking group probably occurred due to lever pressing to L1+S1 in this group. In support of this idea, a 3-way mixed ANOVA on lever press responses during phase 2 (Fig. 4A), followed by a simple main effects comparison (Bonferroni adjustment) showed that there was a significant difference in the lever press response to the two compound cues in sign trackers (F=28.499; p<0.001), but not in goal trackers (F=0.045; p=0.834). There was also a significant difference between the two groups in their lever press response to L1+S1 (F=38.563; p<0.001), but not in their lever press response to L2+S2 (F=3.821; p=0.059). These results suggest that only L1+S1 elicited a lever press conditioned response in the sign trackers. None of the compound cues elicited a lever press conditioned response in the goal trackers.

In addition, figure 4A shows an increasing trend in the lever press response to L2+S2 in the sign tracking group across the sessions of phase 2, suggesting that the sign trackers may have been gradually developing a lever press response to L2+S2 during phase 2. However, the effect of session on L2+S2 in the sign tracking group did not reach significance (F=2.435; p=0.057). There was also no effect of session on the lever press response to L1+S1 in the sign tracking group (F=1.030; p=0.418) or on the lever press response of the goal tracking group to L1+S1 (F=0.28; p=0.92) and L2+S2 (F=0.177; p=0.969).

## Discussion

We examined the expression of the Kamin blocking effect in subjects that develop either a goal tracking or sign tracking conditioned response to a lever cue paired with food reward. We found that animals that sign track show the Kamin blocking effect while this effect is attenuated in the goal tracking group. The use of localized cues in our study allowed the development of both sign and goal tracking behavior. Thus our findings extend previous studies by clarifying the differential expression of blocking in sign and goal trackers

In our experimental design, both Kamin blocking and overshadowing of the sound cue S1 by the lever cue L1 can in theory explain greater responding to sound cue S2 than to S1 during the extinction test in the sign tracking group. However, Aggarwal et al. (2020) showed that L1 failed to overshadow S1 in our experimental paradigm. Thus the differences observed between responding to S1 and S2 in the sign tracking group are due to Kamin blocking, and our results imply that the blocking effect is attenuated in goal trackers.

In our analysis, there were 22 rats in the sign tracking group and 14 rats in the goal tracking group, and thus the difference in the presence of a statistical effect in sign trackers but not in goal trackers is subject to issues of differential power. However, the 22 sign trackers belonged to three different experiments in Aggarwal et al. (2020), with 9, 12 and 10 sign trackers, with five being common in the experiments 2 and 3. In all three experiments, the statistical effect with this reduced number of sign trackers was also significant. This suggests that the differences in statistical power are not critical to the significance of the statistical effects in sign trackers.

The current finding is important because the conditions under which the Kamin blocking effect applies still need to be unraveled. Recently, Maes et al. (2016) reported several failures to replicate the Kamin blocking effect. Our finding that the Kamin blocking effect is attenuated in goal trackers provides insight into some of the conditions necessary for blocking to occur, and suggests that Kamin blocking experiments should use only those subjects that develop a sign tracking response during phase 1. Thus, it is essential to use procedures in which sign tracking and goal tracking responses can be distinguished during phase 1, such as using localized cues which support both sign and goal tracking conditioned responses.

Previous investigations into the expression of the blocking effect in sign and goal trackers have produced mixed results. Holland et al., (2014) showed that goal tracking to auditory cues did not block the acquisition of a sign tracking response to the added lever cues during the compound cue training phase. However, in their experiments, the lever cues overshadowed the formation an association between an auditory cue and food reward. Further, the lever cue when added, during the compound cue conditioning phase, to the previously conditioned auditory cue, took conditioned responding away from the auditory cue. Taken together, their findings suggest that previously conditioned auditory cues failed to attenuate conditioning to added lever cues because of the difference in associability of the auditory and lever cues in their paradigm rather than because of an attenuation in the blocking effect.

In contrast, Derman et al., (2018) showed that both sign tracking and goal tracking animals express the Kamin blocking effect. While this seems to contradict the present results, there is an alternate explanation for their findings, if the details of the training procedure they used are taken into account. First, Derman et al. (2018) used diffuse auditory cues to establish the goal tracking conditioned response. However, diffuse cues only allow for the goal tracking conditioned response. Thus, as mentioned in the introduction, when diffuse cues are used to establish goal tracking, the learning principles being followed may be the same as those underlying the development of a sign tracking conditioned response. This may have caused the animals that developed a goal tracking response to diffuse auditory cues to express blocking in their paradigm.

Second, Derman et al. (2018) used two sound cues in phase 1, one paired with reward (S1→R) and the other not paired with reward (S2→X). In phase 2, they compounded the sound cues by adding simultaneous lever cues (S1+L1 and S2+L2). During phase 2, they presented the previously non-reinforced sound cue (S2) 6 times in the absence of reward (S2→X) and presented its compounded form 4 times followed by reward (S2+L2 →R). Similarly, they presented the previously reinforced sound cue 6 times followed by reward (S1→R) and its compounded form 4 times followed by the reward (S1+L1 →R). This training procedure is expected to lead to the formation of a stronger association between L2 and the reward than a training procedure in which non-reinforced sound cue presentations (S2→X) are not given in phase 2. Therefore, it is suggested here that the use of such a training procedure during compound cue training will lead to the formation of a stronger L2-R association relative to the L1-R association even in the absence of phase 1 (no single cue conditioning phase – no blocking should occur), resulting in a larger response to L2 than to L1 during the post-training extinction test. Derman et al., (2018) indeed found that animals that developed a goal tracking conditioned response showed robust conditioned responding to both the added cues during the extinction test but responded significantly more to L2 than to L1. Derman et al. (2018) argued that this difference in responding reflects the Kamin blocking effect.

The alternative interpretation of Derman et al (2018) suggested here is that the difference in conditioned responding to L2 and L1 observed in the extinction test occurred because of the compound cue training procedure they used. Further, this difference in responding to L1 and L2 is predicted even if the animals did not undergo phase 1 of their behavioral paradigm, even though no blocking should occur in the absence of phase 1. The fact that the goal tracking animals responded robustly to both the lever cues in their extinction test leaves open the possibility that their goal trackers did not express the Kamin blocking effect. Our results clarify the ongoing controversy in the expression of the blocking effect by sign and goal trackers by showing that among animals that develop these two contrasting conditioned responses to a localized lever cue, only sign trackers express the blocking effect.

Recent studies indicate that different neural substrates underlie goal tracking and sign tracking behavior. Systemic (Flagel et al., 2011) and local nucleus accumbens core (Saunders & Robinson, 2012) injections of the dopamine receptor antagonist flupenthixol during classical conditioning impaired both the acquisition and performance of the sign tracking response. These findings suggest that both the *acquisition* and *performance* of the sign tracking conditioned response depend on dopamine in the nucleus accumbens core. In contrast, injecting flupenthixol into the nucleus accumbens core did not impair the acquisition or performance of the goal tracking response (Saunders & Robinson, 2012). The performance of the goal tracking response was impaired under systemic flupenthixol injections (Flagel et al., 2011). This suggests that the *acquisition* of the goal tracking response does not depend on dopamine. The *performance* of the goal tracking response although dependent on dopamine, does not depend on dopamine function in the nucleus accumbens core. These differences in the neural substrates mediating sign tracking and goal tracking conditioned responses imply that different learning mechanisms and principles may underlie the acquisition of these two types of conditioned responses (Derman et al., 2018). Our experimental finding of a difference in the expression of the Kamin blocking effect between animals that either goal track or sign track to the paired lever cue lends further support to the suggestion that different learning mechanisms underlie the development of these two conditioned responses.

Relating the dopamine RPE to sign and goal trackers, Flagel et al. (2011) also found that in goal tracking animals, expected rewards evoke a robust dopamine response. We found that Kamin blocking is attenuated in goal trackers. Conversely, sign tracking animals, in which expected rewards evoke a diminished dopamine response (Flagel et al., 2011), expressed the blocking effect. These findings support the hypothesis that the reduction in dopamine response evoked by the reward when it is expected is necessary for the Kamin blocking effect (Sharpe et al., 2017; Steinberg et al., 2013). However, the current finding seemingly contradicts the prediction error based theoretical explanation of the blocking effect, because the conditioned cue develops predictive properties in both sign and goal tracking animals. This is a topic for a more extended theoretical review (Aggarwal and Wickens, submitted).

## Acknowledgements

This work was supported by the Japan Society for the Promotion of Science Research Fellow Grant-in-Aid 17J00868.

## Notes

**Conflict of Interest Statement** The authors declare no competing interest.

### Competing Interest Statement

The authors have declared no competing interest.

